# O antigen biogenesis sensitises *Escherichia coli* K-12 to bile salts, a likely cause for how it lost its O antigen

**DOI:** 10.1101/2023.05.08.539823

**Authors:** Jilong Qin, Yaoqin Hong, Renato Morona, Makrina Totsika

## Abstract

*Escherichia coli* K-12 is a model organism for bacteriology and has served as a workhorse for molecular biology and biochemistry for over a century since its first isolation in 1922. However, *Escherichia coli* K-12 strains are phenotypically devoid of an O antigen (OAg) since early reports in the scientific literature. Recent studies reported the presence of independent mutations that abolish OAg biogenesis in *E. coli* K-12 strains from the same original source, suggesting unknown evolutionary forces have selected for loss of OAg during the early propagation of K-12. Here, we show for the first time that restoration of OAg in *E. coli* K-12 strain MG1655 synergistically sensitises bacteria to vancomycin with bile salts (VBS). Suppressor mutants surviving lethal doses of VBS mostly contained disruptions in OAg biogenesis. We present data supporting a model where the transient presence and accumulation of lipid-carried OAg intermediates in the bacterial periplasm interfere with peptidoglycan synthesis, causing growth defects that are synergistically enhanced by bile salts. Lastly, we demonstrate that continuous bile salt exposure of OAg-producing MG1655 in the laboratory, can recreate a scenario where OAg disruption is selected for. Hence our work provides a likely explanation for the long-held mystery of how *E. coli* K-12 lost its OAg production and opens new avenues for exploring long-standing questions on the intricate network coordinating the synthesis of different cell envelope components in Gram-negative bacteria.

**Significance statement:** *Escherichia coli* K-12 is the most studied microorganism, widely used in laboratories for studying bacteriology and as a tool for molecular biology. The reason why it is devoid of O antigen remains a long-standing question. Our work has uncovered a previously unknown selection pressure of bile salts on bacterial O antigen biogenesis, which provides a plausible scenario for how the early propagation of *E. coli* K-12 strains in bile salt containing media could have led to loss of O antigen in K-12. Our results also suggest that the accumulation of O antigen intermediates in the bacterial periplasm may interfere with bacterial cell wall synthesis, which paves a new research direction into the interplay of different cell envelope component synthesis pathways.

## Introduction

*Escherichia coli* K-12, the most intensively studied microorganism, has served as a model organism for modern bacteriology and as a workhorse for biochemistry, genetics, and molecular biology. The wild-type strain of *E. coli* K-12 was first isolated in 1922 from the stool of a diphtheria patient and was soon after deposited in the Department of Bacteriology at Stanford University, where it was used in the teaching laboratories and maintained in stab cultures (Fig. 1a)^1,2^. The first report of *E. coli* K-12 in the scientific literature was in 1944^3^ where it was acquired from Dr. C. E. Clifton (Dept. of Bacteriology, Stanford University). This culture was taken to Europe and shared with W. Hayes where it was given the name EMG2^4^. The EMG2 strain subsequently gave rise to strain MG1655^5^, which has since been used widely worldwide as a wild-type *E. coli* K-12 strain and provided the first *E. coli* whole genome sequence published in 1997^6^(Fig. 1a). The original *E. coli* K-12 culture shared from the Tatum laboratory was lost in the Lederberg laboratory and was replaced by another subculture of K-12 sourced from the collection of the Department of Bacteriology at Stanford University that was given the name WG1 (Fig. 1a)^2^. After 30 years of maintenance in laboratories, all *E. coli* K-12 strains were reported by serological studies to be devoid of antigenic structures (O and K antigens) typically found on newly isolated wild-type stains^7^ and were also shown to fail to colonise the human gut^8^.

**Fig 1.**
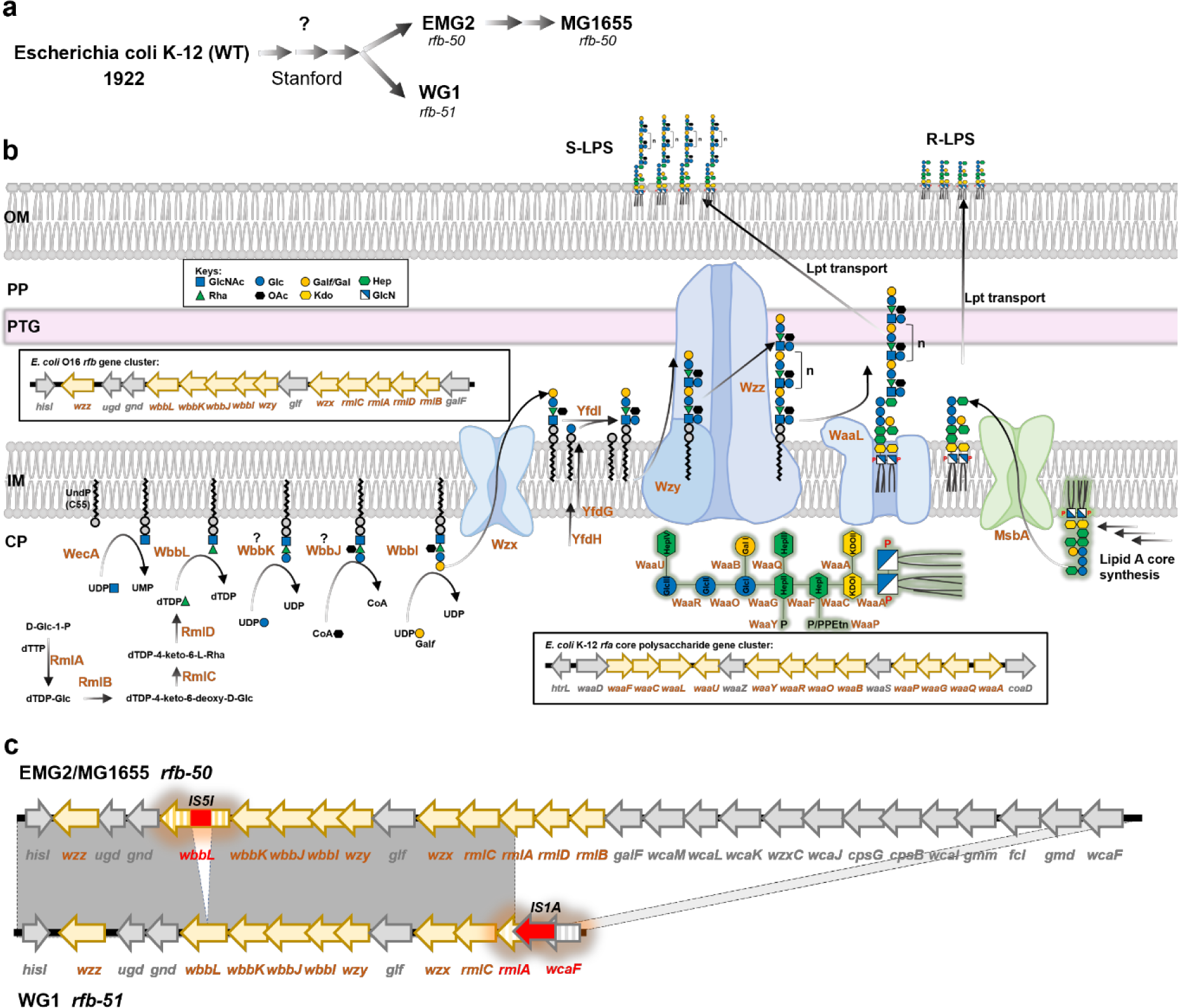
The biogenesis of O16-LPS in *E. coli* K-12. **a)** Schematic representation of the early history of *E. coli* K-12 wild-type strains and derivatives, denoting relevant *rfb* mutations. Details of the procedures and media used during isolation and early propagation of strains are unknown (marked by question mark) but can be inferred by common practices at the time. **b)** Biogenesis of O16-LPS in the original WT *E. coli* K-12 strain. The enzymatic order of glycosyltransferases WbbK and WbbJ is marked with question marks as it remains to be experimentally determined and the order presented is predicted from the inverted gene order within the operon. The structure of Lipid A-core is shown in green glow with corresponding glycosyltransferases and kinases. Genes that are responsible for the biosynthesis of C55-PP-OAg^O16^ and K-12 lipid A-core are shown in box. S-LPS, smooth LPS; R-LPS rough LPS. OM, outer membrane; PP periplasm; PTG, peptidoglycan; IM, inner membrane; CP, cytoplasm. **c)** Schematic representation of sequence alignment between regions *rfb-50* from EMG2/MG1655 and *rfb-51* from WG1. Grey shading marks areas of 100% sequence identity.

The O antigen (OAg) is a major component of bacterial surface lipopolysaccharides (LPS) and in *E. coli* is responsible both for serological specificity, due to its highly antigenically variability^9^, and for host gut colonisation^10^. Presentation of OAg substituted LPS (termed smooth LPS, S-LPS) on the bacterial cell surface requires synthesis of complete core oligosaccharides (by the *rfa* gene cluster) on lipid A, as well as synthesis of OAg repeating units (RUs) (by the *rfb* gene cluster). *E. coli* K-12 was shown to synthesise the complete core oligosaccharides structure^11^ (termed rough LPS, R-LPS) using glycosyltransferases and kinases encoded by *waa* genes in the *rfa* region (Fig. 1b), and was able to be substituted with other OAg when carrying cloned *rfb* gene clusters from other strains^12,13^, suggesting that *E. coli* K-12 is defective in its chromosomal *rfb* region. The K-12 *rfb* region contains genes encoding dTDP-L-rhamnose synthesis enzymes (RmlABCD) and glycosyltransferases for O16 OAg (OAg^O16^) ^14,15^ (Fig. 1b). The synthesis of OAg^O16^ is on a universal lipid carrier undecaprenol phosphate (C55-P), on which the addition of monosaccharides and chemical modifications is sequentially catalysed by WecA, WbbL, WbbK, WbbJ and WbbI to produce the complete O16 repeating unit (RU^O16^) in the cytosolic face of the inner membrane (IM)^16^ (Fig. 1b). The complete unit of C55-PP-RU^O16^ is then flipped across the IM by the Wzx flippase to the periplasmic face of the IM and modified by glucosyltransferase YfdI, the polysaccharide polymerase Wzy and the co-polymerase Wzz (Fig. 1b) and finally WaaL in a competing manner ^17^.

K-12 strains EMG2 and WG1 (both sourced from Stanford University) were found to carry mutations in the OAg gene cluster (*rfb*)^14^ that led to production of R-LPS in both strains (Fig. 1b). Despite their common phenotype, independent mutations were mapped in *rfb*, with EMG2 carrying the *rfb-50* mutation and WG1 carrying the *rfb-51* mutation^14^. The *rfb-50* mutation in strain EMG2 is due to an IS5I insertion element in the coding sequence of WbbL, the second glycosyltransferase of RU^O16^, abolishing the production of OAg^O16^. Mutation *rfb-51* in strain WG1 involves a large deletion between the dTDP-L-rhamnose synthesis gene *rmlA* and the colanic acid synthesis gene *wcaF* (Fig. 1c)^18^. A plasmid carrying the intact *wbbL* from WG1 restored production of S-LPS in the EMG2 strain^14^, confirming the independency of *rfb* mutational events in these two K-12 strains originating from the same institute. This evidence strongly suggests that *E. coli* K-12 was once producing an OAg (i.e. had S-LPS). However, all K-12 strains appear to lack OAg since their first report in literature^3^, suggesting that loss of OAg occurred early in the original isolate’s history, possibly during strain maintenance at Stanford University from the 1920s to 1940s. It is likely that certain culture conditions used at the time for the early propagation of *E. coli* K-12 strains at Stanford University were unfavourable to OAg production, and led to selection of two independent *rfb* mutations in strains WG1 and EMG2.

Isolation and identification of *E. coli* strains in the early 20^th^ century relied on specialised culture media, including MacConkey medium^19^, which through the means of bile salts and crystal violet offered some selectivity by inhibiting the growth of Gram-positive bacteria^20^. MacConkey medium has in fact been used widely among bacteriologists, teachers and practitioners since 1922^21^, as well as in clinical laboratories to identify bacteria from patient stool samples^22,23^. Here, we explored experimentally the selective stress that MacConkey medium could have posed to *E. coli* K-12 for loss of its OAg production. We report that restoration of OAg production in *E. coli* K-12 sensitised bacteria towards vancomycin when grown on MacConkey agar due to the presence of bile salt (BS). We experimentally demonstrate that combining vancomycin with bile salts (VBS) selects for loss of OAg in *E. coli* K-12 strains genetically restored to produce smooth O16 LPS, and that the presence of the C55-PP-OAg intermediate in the periplasm during production of O16 OAg is what sensitised bacteria to BS. Lastly, we demonstrate that culturing smooth *E. coli* K-12 in media supplemented with BS rapidly selects for R-LPS mutants. Taken together our data provide by far the most plausible explanation to the long-standing mystery of how *E. coli* K-12 lost its OAg during early strain maintenance in the laboratory and shed new light into the interplay between different cell envelope component synthesis pathways in Gram-negative bacteria.

## Results

### Restoration of OAg production sensitises E. coli K-12 to vancomycin in the presence of bile salts

To investigate the cause of the loss of OAg in *E. coli* K-12, we firstly restored production of OAg^O16^ in the K-12 reference strain MG1655. An isogenic MG1655-S (‘-S’ for smooth) variant was engineered by flawless allelic exchange with an intact *wbbL* PCR amplicon (Fig. s1 a-b) followed by selection with Colicin E2. Strain MG1655-S was genotypically confirmed by whole genome sequencing and was phenotypically confirmed to produce OAg^O16^ that conferred resistance to Colicin E2 (Fig. s1 c-e).

To examine the potential envelope stress posed by the commonly used MacConkey selective media to both MG1655 and MG1655-S, we supplemented MacConkey Agar with the antibiotic vancomycin, a drug non-permeable for the intact *E. coli* outer membrane. While MG1655 exhibited vancomycin resistance on MacConkey agar as expected, MG1655-S showed a dramatic increase in vancomycin susceptibility under these conditions (Fig. 2a). The selective growth inhibition of most Gram-positive bacteria by MacConkey media is attributed to the presence of crystal violet and BS. However, supplementation of LB agar with BS alone (as sodium deoxycholate, DOC) (Fig. 2b), but not crystal violet (Fig. s2a) in a concentration that showed growth inhibition of a *tolC* mutant of MG1655 (Fig. 2b & Fig. s2a), was sufficient to increase the susceptibility of MG1655-S towards vancomycin. Strikingly, despite MG1655-S exhibiting increased susceptibility to vancomycin in the presence of DOC compared to MG1655 (Fig. 2c), no differences were detected in neither BS resistance (Fig. s2b) nor outer membrane leakage caused by DOC (Fig. s2c) between the strains. Nevertheless, these data suggest that the production of OAg^O16^ in MG1655-S sensitised it to vancomycin in the presence of BS. Indeed, the expression of WbbL *in trans* in MG1655, which restores the production of OAg^O16^, in the presence of DOC and vancomycin resulted in bacterial lysis when cultured (Fig. 2d). In addition, bacterial sensitivity to a combination of DOC and vancomycin (DV) in the presence of OAg^O16^ was not limited to strain MG1655, since OAg^O16^ restoration in K-12 strain UT5600 (Fig. s1f) also sensitised bacteria towards DV (Fig. 2b). Furthermore, production of a different OAg type following introduction of the *Shigella flexneri rfb* region into MG1655 also increased susceptibility to DV treatment (Fig. 2e), suggesting that observed effects were not limited to the specific O16 structure and are likely due to OAg biogenesis.

**Fig 2.**
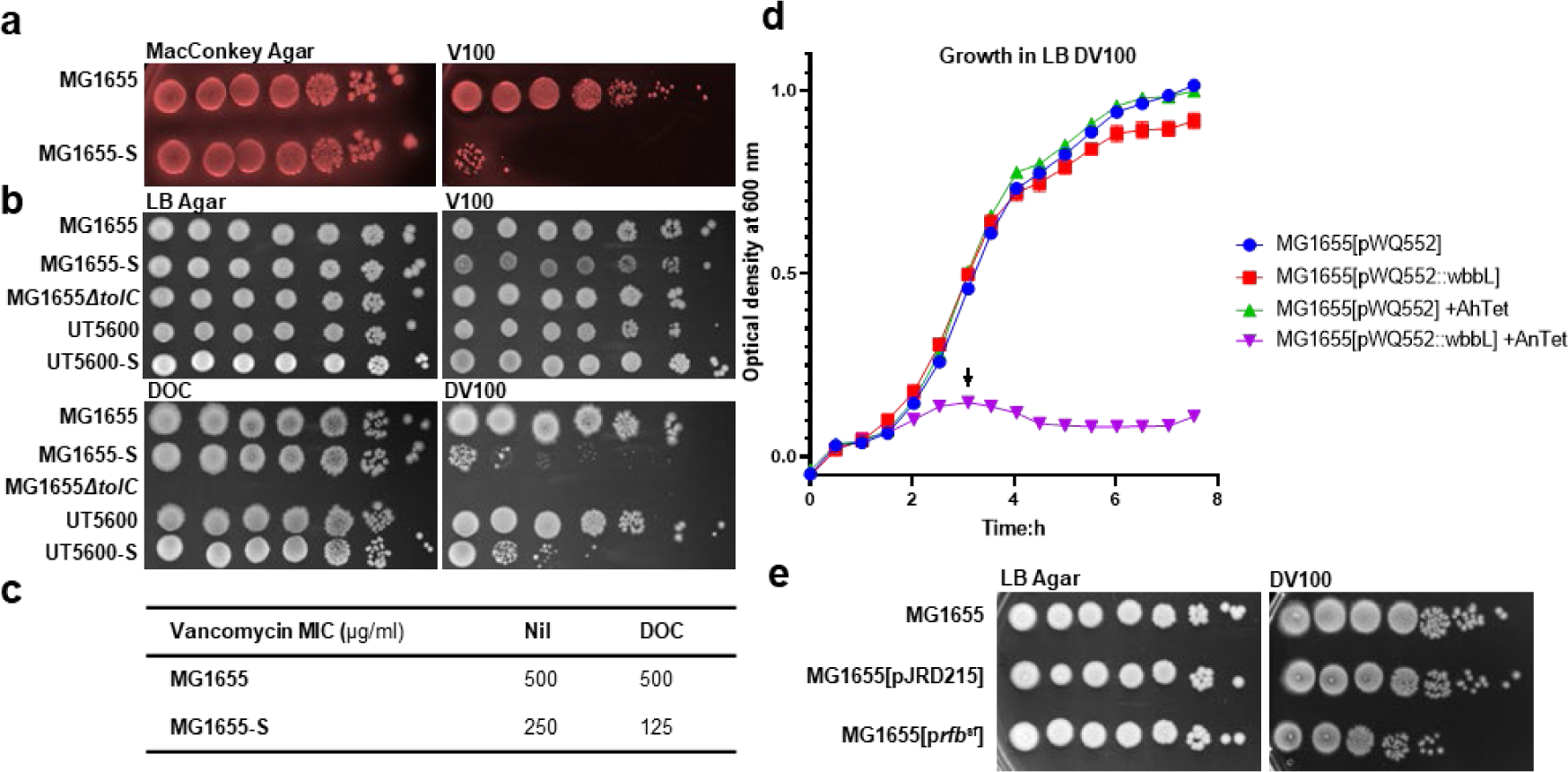
Restoration of O-antigen production sensitises *E. coli* K-12 to bile salts vancomycin. Bacterial cultures of indicated strains grown in LB media were adjusted to OD_600_ of 1 and spotted (4 μl) in 10-fold serial dilutions (10^0^ to 10^-6^) onto MacConkey agar (**a**) or LB agar (**b & e**) supplemented without or with 100 μg/ml vancomycin (V100), 0.1% (w/v) sodium deoxycholate (DOC), or both (DV100). p*rfb*^sf^, pJRD215 carrying *rfb* region from *Shigella flexneri*. **c)** Vancomycin minimum inhibitory concentration (MIC, μg/ml) for indicated *E. coli* strains in the absence (Nil) or presence of DOC. **d)** Growth curves of indicated *E. coli* K-12 strains harbouring plasmids without or with *wbbL* cultured in LB media supplemented with DV100 and with or without 1 ng/ml anhydrotetracycline (AnTet). Arrow indicates lysis.

### Disruption of OAg biogenesis in MG1655-S restores resistance to VBS

To investigate the mechanism of increased susceptibility to VBS in MG1655-S, we collected 51 independent suppressor mutants (BP1-51) recovered following MG1655-S growth on agar plates supplemented with DV. Culturing isolated suppressor mutants in plain LB media exhibited no growth defects compared to MG1655-S or MG1655 (Fig. s3a), yet mutant growth patterns in media supplemented with DV fell into three distinctive groups (Fig. 3): i) early exponential growth followed by lysis similar to MG1655-S parent strain (low DV resistance growth group, termed **L**) (Fig. s3b), ii) intermediate restoration of growth to mid-exponential phase and then stalled (medium DV resistance growth group, termed **M**) (Fig. s3c), and iii) full growth similar to wild-type MG1655 (full DV resistance growth group, termed **F**) (Fig. s3d). Strikingly, LPS profile analysis of suppressor mutants revealed a strong association between S-LPS production and susceptibility to DV (Fig. 3 and Fig. s3e), whereby i) in the **L** group with the least increase in DV resistance, all (27/27) suppressor mutants maintained production of S-LPS, despite some observed alterations in either OAg modal length pattern or band intensity, ii) in the **M** group with intermediate increase in DV resistance, 8 out of 13 suppressor mutants had reduced S-LPS production, and iii) in the **F** group with fully restored resistance to DV, the majority (10/11) of suppressor mutants had undetectable or reduced S-LPS. We also used colicin E2 resistance to uncover any defects in O16-S-LPS that might be undetectable by silver staining. Strikingly, we found that over 60% (32/51) of all suppressor mutants had increased sensitivity to colicin E2 in comparison to the parent strain MG1655-S (Fig. 3). Of those, the majority (24/32) showed medium (**M**) or full (**F**) growth restoration in media supplemented with DV. In contrast, most **L** group suppressors (18/27) exhibited the same colicin E2 resistance as the MG1655-S parent strain (Fig. 3). Together, these data suggest that the production of O16-S-LPS reduces resistance to DV treatment, and that different mechanisms of suppressing VBS stress have been selected in the three groups of isolated MG1655-S suppressors.

**Fig 3.**
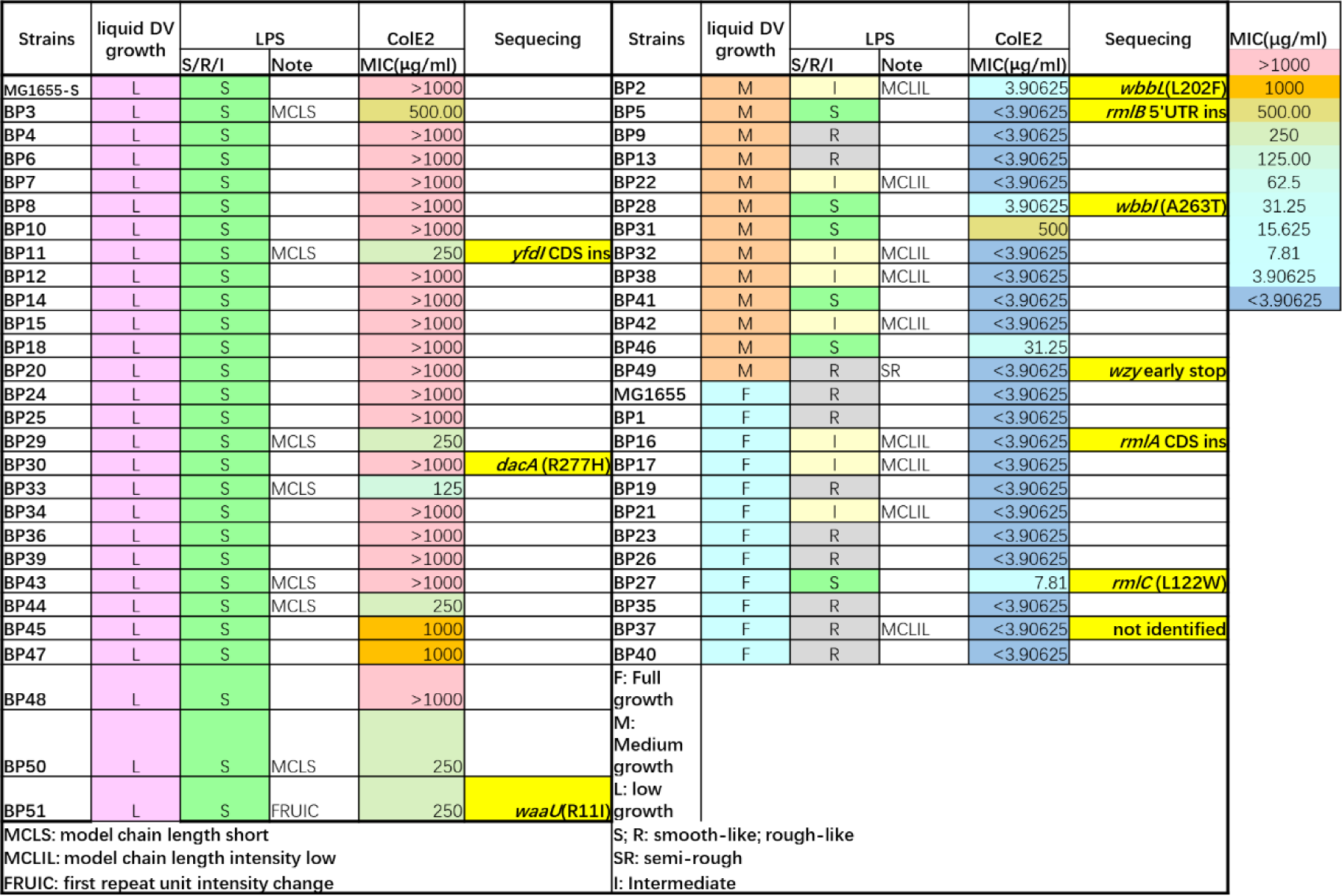
Genotypic and phenotypic traits of isolated MG1655-S suppressor mutants. Growth profiles of 51 MG1655-S suppressor mutants (BP1-51) in LB supplemented with 100 μg/ml vancomycin and 0.1% (w/v) sodium deoxycholate (DV) were classified as low (L) and similar to MG1655-S, medium (M) with growth recovery in between MG1655 and MG1655-S, or full (F) and similar to MG1655. The LPS silver staining profile of each mutant was compared to MG1655-S and MG1655 and classified as smooth-like LPS (S), rough-like LPS (R), or intermediately O antigen substituted LPS (I) with additional detailed alterations described in notes. The MIC values (μg/ml) of purified colicin E2 (ColE) are recorded for each mutant. Suppressor mutants selected for whole genome sequencing are highlighted in yellow shading and their identified mutations are detailed by gene/coding sequence (CDS) name and non-synonymous substitution, early stop or insertion (ins). The mutation in BP49 was identified in *wzy* by targeted Sanger sequencing.

To study the suppressor genotypes, we chose 3 representative mutants from each of the three DV growth recovery groups (**L**, **M**, and **F**) and analysed them by whole genome sequencing (Fig. 3). We found that 7 of the 9 suppressor mutants had mutations in O16-S-LPS synthesis pathways. Specifically, mutations were identified in: the glycosyltransferase *yfdI* gene responsible for glucose modification of OAg^O16^ in the periplasm (BP11); in K-12 LPS outer core synthesis gene *waaU* (BP51); in genes required for the synthesis of OAg^O16^ in the cytoplasm (*wbbL* in BP2 and *wbbI* in BP28); and in genes responsible for dTDP-L-rhamnose synthesis (an OAg^O16^ sugar) (*rmlA* in BP16, *rmlB* in BP2 and *rmlC* in BP27). Suppressor mutant BP49, which showed an LPS profile with only one RU^O16^ substitution (Fig. s3e), was confirmed to have a mutation in gene *wzy* encoding OAg polymerase by targeted sequencing of the *wzy* locus. In addition, in mutant BP30 we identified a mutation in *dacA*, that is a gene responsible for peptidoglycan carboxypeptidase, known as penicillin binding protein 5 (PBP5). Disruption of *dacA* was shown to have increased vancomycin resistance in MG1655 here (Fig. s4a) and previously^24^. However, the isolated DacA^R227H^ mutation conferred the least DV resistance in BP30 (Fig. 3), suggesting that the main stress posed by DV treatment on MG1655 is from the presence of DOC and is synergistically enhanced by vancomycin. Interestingly, we did not identify any genetic changes in the genome of BP37, a suppressor mutant showing full restoration of DV resistance and an altered LPS profile. To validate suppressor mutant findings, we generated single gene deletion mutants of *rmlC* and *wzy* in MG1655-S. The resulting strains showed restored resistance to DV treatment (Fig. 4a), similar to isolated suppressor mutants. In addition, complementation by *rmlC*^WT^ but not *rmlC*^L122W^ (suppressor mutation in BP27) restored the production of S-LPS in MG1655-S*ΔrmlC* (Fig. s4b) and re-sensitised it to DV (Fig. s4c). Ectopic expression of RmlC^WT^ in BP27 also re-sensitised this suppressor mutant to DV (Fig. s4d). These data confirmed that the suppressor mutations identified in genes involved in O16-S-LPS synthesis in MG1655-S are directly responsible for regaining of VBS resistance in these strains. Taken together, our genotypic analyses confirmed the phenotypic finding that biogenesis of O16-S-LPS in MG1655-S sensitised bacteria towards VBS treatment.

**Fig 4.**
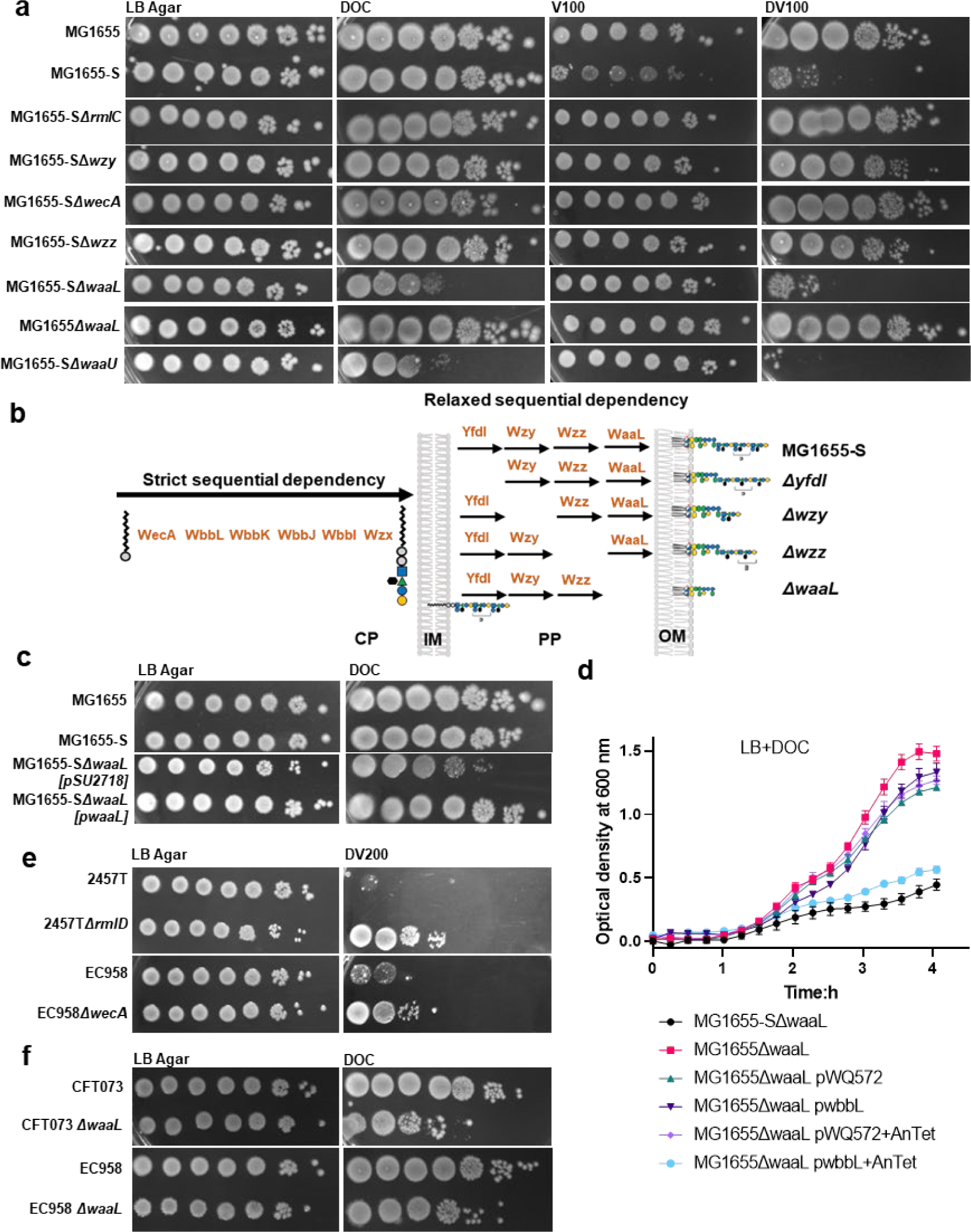
Accumulation of O antigen in the periplasm sensitises *E. coli* K-12 to bile salts. Bacterial cultures of indicated strains grown in LB media were adjusted to OD_600_ of 1 and spotted (4 μl) in 10-fold serial dilutions (10^0^ to 10^-6^) onto LB agar (**a, c, e, f**) without or supplemented with 100 μg/ml vancomycin (V100), 0.1% (v/w) sodium deoxycholate (DOC), a combination of V100 and DOC (DV100) or DOC and 200 μg/ml vancomycin (DV200). (**b**) Schematic representation of the synthesis and assembly process of O16 substituted LPS in different MG1655-S mutants. (**d**) Growth curves of indicated *E. coli* K-12 strains harbouring plasmids without or with *wbbL* cultured in LB media supplemented with 0.1%(w/v) DOC and without or with 1 ng/ml anhydrotetracycline (AnTet).

### Presence of C55-PP-OAg in periplasm sensitises bacteria to bile salts

Disruption of OAg synthesis and/or assembly enhances production of enterobacterial common antigen (ECA) in *E. coli* due to the shared initial substrate C55-PP-GlcNAc that can be utilised both by the OAg^O16^ second glycosyltransferase WbbL and the ECA second glycosyltransferase WecG^25^. In addition, ECA has been reported previously to be important for DOC resistance^26^. However, ECA in MG1655-S mutants disrupted for OAg synthesis and/or assembly did not account for their increased resistance to DV since MG1655-S*ΔwecA* that has lost the production of both OAg and ECA (due to inability to synthesise the common precursor C55-PP-GlcNAc) (Fig. s4f) also showed restored DV resistance (Fig. 4a). In addition, we also generated a single deletion of *wzz* mutant of MG1655-S (encodes the OAg co-polymerase responsible for modal chain length control). Interestingly, disruption of *wzz*, also alleviated DV susceptibility (Fig. 4a). Taken together, the inactivation of genes that catalyse processing steps of OAg^O16^ in the periplasm, such as *yfdI*, *wzy* and *wzz* (Fig. 4a) restored resistance to DV, similar to disrupting genes responsible for OAg^O16^ synthesis in the cytosol (*rmlA, rmlB, rmlC, wbbL, wbbI, wecA*) (Fig. 3 & Fig. 4a).

Synthesis of OAg^O16^ in the cytosol is strictly sequential (Fig. 4b) as it depends on the distinct substrate specificity of each of the various glycosyltransferases. Synthesis of RU^O16^ will stall when either a glycosyltransferase or its sugar substrate is missing. On the contrary, the modification of OAg^O16^ (by YfdI), its polymerisation (by Wzy/Wzz) and ligation (by WaaL) in the periplasm have relaxed sequential dependency (Fig. 4b) due to competition over universal substrates, where the absence of one does not impede the function of the rest. It is possible that disrupting the genes responsible for OAg cytosolic synthesis in MG1655-S completely abolished or delayed the accumulation of C55-PP-OAg^O16^ into the periplasm, while deletion of genes responsible for C55-PP-OAg^O16^ periplasmic processing accelerated its clearance from the periplasm (through translocation across the outer membrane by the Lpt apparatus^27^). To test this hypothesis, we generated the ligase mutant MG1655-S*ΔwaaL* (Fig. s4f) that is unable to transfer OAg^O16^ from C55-PP onto the lipid A core and would thus accumulate C55-PP-OAg^O16^ in the periplasm (Fig. 4b)^28^. While disruption of *waaL* in MG1655 did not affect resistance to DV, in MG1655-S disruption of *waaL* failed to restore resistance to DV (Fig. 4a) despite affecting LPS production (the mutant has a R-LPS profile, Fig. s4f). Indeed, a lipid A core mutant MG1655-S*ΔwaaU* that is unable to accept OAg^O16^ (Fig. s4f) and would also accumulate C55-PP-O16 in the periplasm also failed to restore resistance to DV (Fig. 4a). These findings suggest that accumulation of C55-PP-OAg^O16^ in the periplasm sensitised the strain to DV. Intriguingly, susceptibility was due to increased sensitivity to DOC (Fig. 4a&c) as both MG1655-S*ΔwaaL* and MG1655S*ΔwaaU* unexpectedly exhibited increased vancomycin resistance (Fig. s4e). Similarly, restoration of OAg^O16^ production by WbbL expression in MG1655*ΔwaaL* slowed down its growth in the presence of DOC to the same level as MG1655-S*ΔwaaL* (Fig. 4d), confirming that DOC poses stress on the production of C55-PP-OAg^O16^ in the periplasm. Collectively, these data suggest a model where either accumulation or delayed clearance of C55-PP-OAg^O16^ in the periplasm sensitises MG1655-S towards BS.

### BS sensitisation to OAg biogenesis exists among other S-LPS producing strains

Since *E. coli* K-12 has been heavily domesticated in the laboratory, we questioned whether sensitivity to DV can also be observed in other strains, including clinically relevant isolates. Surprisingly, we found that *S. flexneri* 2457T*ΔrmlD* with disrupted OAg rhamnose saccharide synthesis, and uropathogenic *E. coli* (UPEC) ST131 EC958*ΔwecA* with disrupted OAg and ECA synthesis all showed increased resistance to DV compared to the wild-type parent strains (Fig. 4e). In addition, deletion of *waaL* in UPEC clinical isolates EC958 and CFT073 sensitised them to DOC (Fig. 4f). These data support that the effect observed in MG1655-S, where presence of C55-PP-OAg sensitises bacteria towards BS, is likely shared widely by other *E. coli* smooth strains and presumably can be extended to other S-LPS producing Enterobacteriaceae.

### Prolonged exposure of MG1655-S to BS promotes loss of OAg

Our results showed that disruption of OAg biogenesis in *E. coli* K-12 increased resistance to VBS, and that the stress is linked to the presence of C55-PP-OAg in the periplasm and BS. This suggests that loss of OAg in K-12 could be an adaptation to culture on BS-containing media. To test this tenet, we carried out an experimental evolution experiment where we continuously cultured MG1655-S in the presence or absence of DOC for up to 28 days and then challenged the cultures with both DV and colicin E2. At day 1 cultures showed no difference in DV resistance or colicin E2 susceptibility (Fig. 5). By day 7, DOC-treated MG1655-S showed a 10-fold increase in DV resistance and a 10-fold decrease in colicin E2 resistance compared to untreated MG1655-S (Fig. 5), strongly indicating that OAg biogenesis was disrupted in DOC-treated cells. Differences were further increased up to 1000-fold at day 28 (Fig. 5), suggesting that prolonged exposure of MG1655-S to BS can positively select for disruption of OAg synthesis.

**Fig 5.**
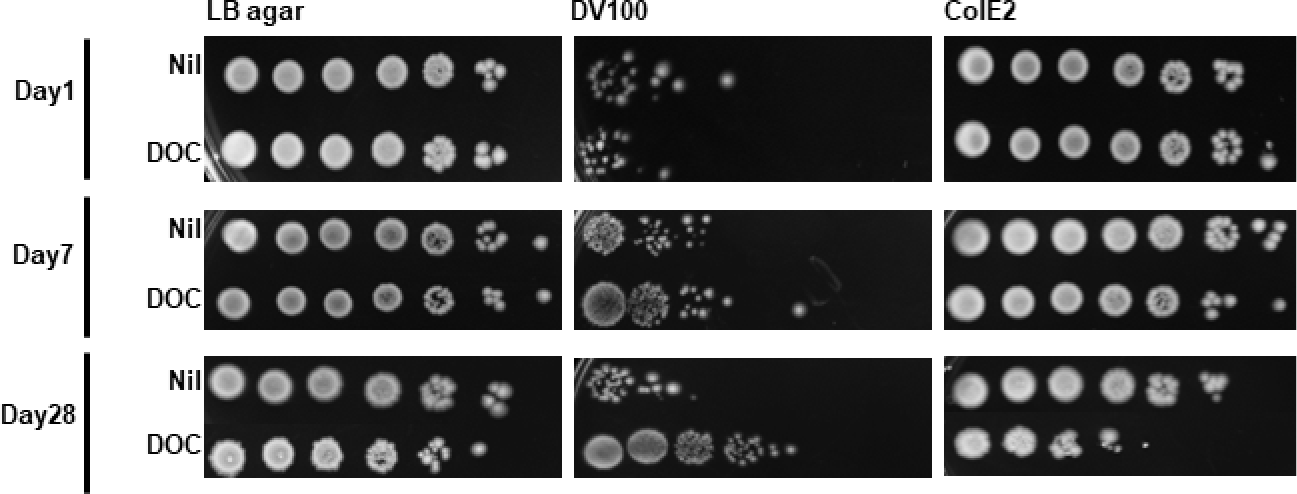
Continuous exposure of MG1655-S to bile salts promotes loss of O antigen. Bacterial cultures of MG1655-S grown in LB media supplemented without (Nil) or with 0.1% (w/v) deoxycholate (DOC) for 1 day, 7 days or 28 days were adjusted to OD_600_ of 1, and spotted (4 μl) in 10-fold serial dilutions (10^0^ to 10^-6^) onto LB agar without or with 100 μg/ml vancomycin and 0.1% (w/v) sodium deoxycholate (DV100), or 20 μg/ml crude colicin E2 preparation (ColE2). Data represent at least three independent repeats.

## Discussion

The surface exposed lipopolysaccharide OAg is important for *E. coli* colonisation of host digestive tracts^8,29,30^. The OAg forms a protective surface layer around bacterial cells helping to withstand gut environmental challenge, including from host antimicrobial peptides (AMPs)^31^ and gut residing bacteriophages, such as the R-LPS attacking coliphage T4^32,33^. Originally isolated from human stools, *E. coli* K-12 strains however were found to be sensitive to coliphage T4 and unable to colonise the human gut^8^. This together with subsequent genetic evidence of independent inactivation of OAg^O16^ biogenesis in two *E. coli* K-12 strains originating from the same institute, strongly suggested that *E. coli* K-12 produced S-LPS upon isolation but was subjected to unknown selection pressure leading to early loss of OAg production. Here we investigate the potential reason for the loss of OAg among *E. coli* K-12 strains by focusing on its isolation and early maintenance in the 1920s. Biles salts, which are produced in the liver and released into the intestine, have been employed since the 1890s in bacterial culture as a key ingredient in selective media for the isolation and culturing of bacteria from human and animal faeces^20^. Indeed, *E. coli* can tolerate high concentrations of BS mainly due to the presence of an outer membrane that limits BS influx into cells, and by actively effluxing intracellular BS through the AcrAB-TolC pump^34^. Although BS are known to cause damage to membranes^35^, DNA^36^, and proteins^37^, the exact mechanism of BS susceptibility in Gram-positive and Gram-negative bacteria with defective efflux pumps^38^ still remains unclear.

Here we showed that while production of OAg in *E. coli* K-12 did not affect resistance to BS, it increased sensitivity to VBS. Since BS treatment indistinguishably caused outer membrane leakage in both MG1655 and MG1655-S, the increased sensitivity to vancomycin observed in MG1655-S is therefore likely not a direct result of outer membrane disruption. Indeed, in the absence of BS, MG1655-S exhibited a 2-fold increase in sensitivity to vancomycin in comparison to MG1655. Vancomycin effects are well studied in Gram-positive bacteria, where it’s been shown to bind to D-Ala-D-Ala at the terminus of pentapeptide linked peptidoglycan (PTG)^39^ and inhibit the transpeptidation of PTG catalysed by penicillin binding proteins (PBPs). Our results in *E. coli* K-12 hence suggest a complex interplay may exist between biogenesis of OAg and PTG in Gram-negative bacteria, where the production of OAg potentially interferes with the biosynthesis and/or maintenance of PTG, leading to *E. coli* K-12’s altered sensitivity to the limited amount of vancomycin that penetrates across the outer membrane. This hypothesis is supported by the fact that the synthesis of both PTG and OAg requires the universal lipid carrier C55-P, thus an interference likely exists between the two pathways through competition for this common lipid carrier. Moreover, the polymerisation of PTG precursor (lipid II) generates C55-PP in the periplasm, which was reported as a potent inhibitor of the PBP1b (MrcB/PonB) in *E. coli* K-12^40^. The production of OAg also generates C55-PP in the periplasm, both by the polymerisation (Wzy) and the ligation (WaaL) process^41^. It is likely, therefore, that PBP1b activity is altered when OAg production is restored in *E. coli* K-12, resulting in increased sensitivity to vancomycin.

We also observed a synergistic effect between BS and vancomycin in inhibiting the growth of MG1655-S but not MG1655. Synergistic effects were also seen in UPEC and *S. flexneri* wild-type strains (naturally producing S-LPS) but not in isogenic mutants with disrupted OAg synthesis, suggesting that our observations in *E. coli* K-12 are true more widely among the Enterobacteriaceae. Synergy of BS and vancomycin was also previously reported in other *E. coli* strains and an *Acinetobacter lwoffi* strain^42^. The synergy mechanism between BS and vancomycin against strains producing OAg remains unclear, as the target(s) of BS in *E. coli* remains unknown. Through our MG1655-S suppressor mutant studies, we found that most suppressors had defects in OAg synthesis pathways. Disruption of early OAg synthesis steps in the cytosol (*rmlABCD* and *wbbL*) can be expected to confer resistance to VBS, as it completely abolishes the use of C55-PP-GlcNAc in OAg synthesis and eliminates any potential interference with PTG synthesis. This is exemplified by the *rfb-50* and *rfb-51* mutations in EMG2 and WG1, respectively. However, we also observed increased resistance to VBS among mutants with disrupted YfdI, Wzy and Wzz, which are enzymes responsible for later processing of OAg in the periplasm, and which compete for C55-P-OAg^O16^ with the WaaL ligase. It is possible that competition between these enzymes over C55-PP-OAg^O16^ limited the rate of O16 release from C55-PP by WaaL in the periplasm, resulting in increased bacterial sensitivity to VBS. Indeed, an increase in BS sensitivity was observed in MG1655-S*ΔwaaL*, CFT073*ΔwaaL* and EC958*ΔwaaL*, but not in MG1655*ΔwaaL,* suggesting that the stalled release and accumulation of C55-PP-Oag in the periplasm sensitised the bacteria to BS. This could in turn have affected peptidoglycan synthesis through competition for C55-PP. Interestingly, we have also observed increased vancomycin resistance upon *waaL* disruption in MG1655-S. Disruption of the D,D-carboxypeptidase DacA (PBP5) in MG1655 was previously reported to increase vancomycin resistance^24^ and here in MG1655-S in the presence of BS. Whether the presence of C55-PP-OAg in the periplasm affects the activity of PBPs remains to be determined in future work currently underway. Importantly, the current study lays the foundation for exploring a new layer of interplay between OAg and peptidoglycan synthesis pathways. Based on our findings, we propose a model for explaining how *E. coli* K-12 lost its OAg, whereby the production of OAg generates transient C55-PP-OAg intermediate in the periplasm, and upon prolonged exposure to BS this enhances interference with PTG synthesis pathways and growth inhibition which can be alleviated by disruption of OAg synthesis.

Regardless of the complex action of VBS in selecting MG1655-S mutants with disrupted OAg biogenesis, we were able to show that the stress posed by BS on MG1655-S was aided by vancomycin killing. A common way of preserving bacterial strains in the past was through stab agar culture (master culture) maintained at room temperature^4^. This practice potentially accelerated strain in-lab adaptation due to starvation in these long-term cultures^43^. However, here we showed that static culture of a genetically restored OAg^O16^ producing *E. coli* K-12 (MG1655-S) at room temperature for as long as 28 days did not select for differences in sensitivity to VBS or colicin E2, suggesting that in-lab starvation alone could not pose adequate selection pressure for MG1655-S to lose its OAg production. Only when we cultured MG1655-S in commonly used media containing BS, could we select for characteristics of OAg biosynthesis disruption in over 90% of the culture population only within 1 week. It is likely that the selection pressure posed on MG1655-S in our in-lab adaptation experiments was a combination of the effects posed by BS on the transient presence of C55-PP-OAg in the periplasm discussed above, and the oxidative stress of exposure to long-term rich media^44^.

Our findings clearly demonstrate a role for BS in disrupting OAg production in *E. coli* K-12 and other Enterobacteriaceae and would argue against routine use of MacConkey and other BS-containing media for prolonged cultures in bacteriology. Their value lies as secondary indicative media in aiding the identification of Enterobacteriaceae from faecal samples. While we are not in a position to trace the exact history of isolation, maintenance and manipulation of the original *E. coli* K-12 strain, or whether BS was definitively used during the passages of *E. coli* K-12 cultures for strain purification, our study provides for the first time the most likely explanation of how the original *E. coli* K-12 lost its OAg.

## Materials and Methods

### Bacterial strains, plasmids and growth media

The bacterial strains and plasmids used in this work are listed in Table s1. Single colonies of *E. coli* strains grown overnight on Lennox Broth (LB) (Bacto Tryptone (#211699, Gibco) 10g/L, Bacto Yeast Extract (#212750, Gibco) 5g/L, NaCl 5 g/L)^45^ agar (1.5% w/v) plates were picked and grown overnight in LB at 37 °C for all experiments. Other bacterial growth media used in this study were Mueller Hinton (#275730, BD), MacConkey Agar (#CM0115, Oxiod). Where appropriate, media were supplemented with ampicillin (Amp, 100 µg/ml), kanamycin (Kan, 50 µg/ml), chloramphenicol (Chl, 25 µg/ml), bile salts sodium deoxycholate (DOC, 0.1% w/v, #D6750, Sigma), vancomycin (Van, 100 µg/ml, #SBR00001, Sigma), L-arabinose (0.2% w/v), D-glucose (0.2% w/v), crystal violet (0.0005% w/v), and colicin E2 (20 µg/ml).

### Site-directed mutagenesis via allelic exchange

*E. coli* gene inactivation mutants were generated by Lambda Red mutagenesis as described previously^46^ with the oligos listed in Table S1. For restoration of *wbbL* in K-12 strains MG1655 and UT5600, an amplicon containing the undisrupted WbbL coding sequence from pPR2191^47^ was generated by PCR with the oligos listed in Table s1, and was then used for allelic exchange gene replacement as above. Successful revertants were selected on LB agar supplemented with 20 µg/ml purified colicin E2 protein. Removal of the IS5I elements in the original *wbbL* gene was confirmed via PCR and phenotypic restoration of O16 substituted LPS was confirmed with SDS-PAGE LPS silver staining (see SI). Whole genome sequencing of the resulting MG1655-S strain by DNBSEQ confirmed the successful removal of IS5I in *wbbL* without the introduction of any other genetic modifications.

### Plasmid construction

The RmlC expression constructs used in this study were constructed by PCR amplification of the *rmlC* alleles from strains MG1655 (*rmlC*^WT^) and BP27 (*rmlC*^L122W^) using the Q5 high-fidelity polymerase (NEB) and oligos listed in Table s1 and subsequent insertion into pBAD18-Chl via restriction digestion cloning.

### Suppressor mutant selection

Suppressor mutant selection of MG1655-S to vancomycin and deoxycholate combination treatment was done by spreading an overnight MG1655-S culture grown in LB (1 in 2000 dilution in fresh LB media) onto LB agar supplemented with 100 µg/ml vancomycin and 0.1% (w/v) sodium deoxycholate, and subsequent incubation at 37 °C overnight. Colonies were carefully streaked out onto non-selective LB plates and stored at −80 °C for downstream analyses.

### Whole Genome Sequencing

For bacterial whole genome sequencing, genomic DNA of MG1655, MG1655-S and suppressor mutants was prepared using a Qiagen QIAamp DNA blood mini kit according to the manufacturer’s protocol. Samples were prepared for DNBseq DNA library construction (BGI) followed by DNBSEQ PE150 sequencing (BGI). The processed reads for each strain were then mapped onto the NCBI MG1655 reference genome (Accession number U00096) using Geneious 8.0. Sequence changes between our laboratory MG1655 and the online reference genome, as well as within-population variations were determined using the Geneious built-in function and are excluded from the analysis in MG1655-S and its suppressor mutants (Table s2).

### Vancomycin and bile salts susceptibility assays

For vancomycin susceptibility testing, overnight cultures of bacterial strains grown in LB were adjusted to OD_600_ of 1 and spotted (4 µl) in 10-fold serial dilutions onto MacConkey agar supplemented with or without vancomycin at different concentrations. Culture dilutions were also spotted onto LB agar supplemented without or with vancomycin, deoxycholate, and crystal violet at different concentrations, as indicated in figure legends. Plates were incubated overnight at 37 °C and then imaged.

### Colicin E2 purification and susceptibility assay

Colicin E2 protein was purified and used to determine the susceptibility of bacterial strains as described previously^48^.

### Bacterial Growth assays

Growth curves for bacterial strains were generated as described previously^49^. Briefly, bacterial strains grown overnight in LB were diluted 1 in 1000 in 200 µl of LB in flat-bottom 96 well plates supplemented without or with 0.1% (w/v) sodium deoxycholate and/or 100 µg/ml vancomycin. For expression of plasmid-borne *wbbL* under *tet* promoter control, 1 ng/ml anhydrotetracycline was also supplemented in media. Microtiter plates were incubated at 37 °C with aeration in a CLARIOstar plate reader (BMG, Australia) programmed to measure the absorbance (O.D. 600 nm) every 6 minutes over 18 h.

### Antibiotic susceptibility testing

Vancomycin minimum inhibitory concentrations (MIC) were determined for bacterial strains according to the Clinical and Laboratory Standards Institute guideline^50^. Briefly, overnight bacterial cultures in LB were sub-cultured in MH broth to a McFarland turbidity standard of 0.5. Cultures were then adjusted to 10^7^ cells/ml and 10 µl samples were spotted onto MH agar plates supplemented with vancomycin at concentrations from 500 µg/ml to 7.8 µg/ml in 2-fold serial dilutions in the presence or absence of 0.1% (w/v) sodium deoxycholate. Plates were incubated overnight at 37°C. The MIC value was determined as the lowest concentration of vancomycin at which bacterial lawn growth was inhibited.

### In vitro adaptation to bile salts

MG1655-S grown overnight at 37 °C in LB broth was sub-cultured 1:1000 in 3 ml LB broth in a 15 ml tube supplemented with or without 0.1% (w/v) sodium deoxycholate in triplicate. Tube lids were loosely attached to allow aeration and tubes were incubated at 21 °C statically for 1 day, 7 days and 28 days. Cultures from different time-points were 10-fold serially diluted and spotted onto LB agar plates supplemented with or without 100 µg/ml vancomycin and 0.1% (w/v) sodium deoxycholate or 20 µg/ml colicin E2, and plates were incubated overnight at 37 °C and imaged. For quantification, cultures were diluted and spread onto LB plates and incubated overnight at 37 °C to isolate single colonies. One hundred colonies from each culture were patched on LB agar plates supplemented with or without 100 µg/ml vancomycin and 0.1% (w/v) deoxycholate or 20 µg/ml colicin E2, and plates were incubated overnight at 37 °C and imaged.

## Acknowledgement

Authors acknowledge funding support from the Australian Research Council (DP210101317), the Max Planck Queensland Centre and a Georgina Sweet Award for Women in Quantitative Biomedical Science to MT. The Ian Potter Foundation sponsored the CLARIOStar high-performance microplate reader (BMG, Australia).

## Author contributions

JQ contributed to project conception and design, conducted experiments, and contributed to data collection, analysis, and interpretation and model proposal; MT supervised the study, contributed to data interpretation and obtained the funding, substantially revised the manuscript; YH contributed to the strain construction and data interpretation; RM contributed to data interpretation and funding. JQ drafted the manuscript, all authors contributed to the manuscript editing.

## Data Availability

All data generated or analysed during this study are included in this published article (and its Supplementary Information files).

## Additional Information

The authors declare no competing interests.

## References

1 Tatum, E. L. & Lederberg, J. Gene Recombination in the Bacterium Escherichia coli. J Bacteriol 53, 673–684, doi:10.1128/jb.53.6.673-684.1947 (1947).

2 Bachmann, b. J. in Escherichia coli and Salmonella: Cellular and Molecular Biology Vol. 2 (ed Frederick C. Neidhardt) Ch. F, (ASM Press, 1996).

3 Gray, C. H. & Tatum, E. L. X-Ray Induced Growth Factor Requirements in Bacteria. Proceedings of the National Academy of Sciences of the United States of America 30, 404–410, doi:10.1073/pnas.30.12.404 (1944).

4 R. C. Clowes., W. H. Experiments in microbial genetics. 194 (John Wiley & Sons Inc, 1968).

5 Guyer, M. S., Reed, R. R., Steitz, J. A. & Low, K. B. Identification of a sex-factor-affinity site in E. coli as gamma delta. Cold Spring Harb Symp Quant Biol 45 **Pt** **1**, 135–140, doi:10.1101/sqb.1981.045.01.022 (1981).

6 Blattner, F. R. et al. The complete genome sequence of Escherichia coli K-12. Science 277, 1453–1462, doi:10.1126/science.277.5331.1453 (1997).

7 Orskov, F. & Orskov, I. The fertility of Escherichia coli antigen test strains in crosses with K 12. Acta Pathol Microbiol Scand 51, 280–290 (1961).

8 Smith, H. W. Survival of orally administered E. coli K 12 in alimentary tract of man. Nature 255, 500–502, doi:10.1038/255500a0 (1975).

9 Kauffmann, F. Studies on the serology of the Escherichia coli group. J Bacteriol 51, 126 (1946).

10 Ho, T. D. & Waldor, M. K. Enterohemorrhagic Escherichia coli O157:H7 gal mutants are sensitive to bacteriophage P1 and defective in intestinal colonization. Infect Immun 75, 1661–1666, doi:10.1128/IAI.01342-06 (2007).

11 Prehm, P., Schmidt, G., Jann, B. & Jann, K. The cell-wall lipopolysaccharide of Escherichia coli K-12. Structure and acceptor site for O-antigen and other substituents. Eur J Biochem 70, 171–177, doi:10.1111/j.1432-1033.1976.tb10967.x (1976).

12 Schmidt, G. Genetical studies on the lipopolysaccharide structure of Escherichia coli K12. J Gen Microbiol 77, 151–160, doi:10.1099/00221287-77-1-151 (1973).

13 Cheah, K. C., Beger, D. W. & Manning, P. A. Molecular cloning and genetic analysis of the rfb region from Shigella flexneri type 6 in Escherichia coli K-12. FEMS Microbiol Lett 67, 213–218, doi:10.1016/0378-1097(91)90356-f (1991).

14 Liu, D. & Reeves, P. R. Escherichia coli K12 regains its O antigen. Microbiology (Reading) 140 (Pt 1), 49–57, doi:10.1099/13500872-140-1-49 (1994).

15 Stevenson, G. et al. Structure of the O antigen of Escherichia coli K-12 and the sequence of its rfb gene cluster. J Bacteriol 176, 4144–4156, doi:10.1128/jb.176.13.4144-4156.1994 (1994).

16 Reeves, P. R. & Cunneen, M. M. in Microbial Glycobiology (eds Otto Holst, Patrick J. Brennan, Mark von Itzstein, & Anthony P. Moran) 319-335 (Academic Press, 2010).

17 Hong, Y. et al. Repeat-Unit Elongations To Produce Bacterial Complex Long Polysaccharide Chains, an O-Antigen Perspective. EcoSal Plus, eesp00202022, doi:10.1128/ecosalplus.esp-0020-2022 (2023).

18 Browning, D. F., Hobman, J. L. & Busby, S. J. W. Laboratory strains of Escherichia coli K-12: things are seldom what they seem. Microb Genom 9, doi:10.1099/mgen.0.000922 (2023).

19 MacConkey, A. T. Note on a New Medium for the Growth and Differentiation of the Bacillus Coli Communis and the Bacillus Typhi Abdominalis. The lancet 156, 20, doi:https://doi.org/10.1016/S0140-6736(01)99513-3 (1900).

20 Macconkey, A. T. Bile Salt Media and their advantages in some Bacteriological Examinations. J Hyg (Lond) 8, 322–334, doi:10.1017/s0022172400003375 (1908).

21 Heinemann, P. G. A laboratory guide in bacteriology: for the use of students, teachers and practitioners. 4 edn, (The Univeristy of Chicago Press, 1922).

22 Jung, B. & Hoilat, G. J. MacConkey Medium. (StatPearls Publishing, 2022).

23 Macconkey, A. Lactose-Fermenting Bacteria in Faeces. J Hyg (Lond) 5, 333–379, doi:10.1017/s002217240000259x (1905).

24 Park, S. H. et al. Divergent Effects of Peptidoglycan Carboxypeptidase DacA on Intrinsic beta-Lactam and Vancomycin Resistance. Microbiol Spectr 10, e0173422, doi:10.1128/spectrum.01734-22 (2022).

25 Barr, K., Ward, S., Meier-Dieter, U., Mayer, H. & Rick, P. D. Characterization of an Escherichia coli rff mutant defective in transfer of N-acetylmannosaminuronic acid (ManNAcA) from UDP-ManNAcA to a lipid-linked intermediate involved in enterobacterial common antigen synthesis. J Bacteriol 170, 228–233, doi:10.1128/jb.170.1.228-233.1988 (1988).

26 Ramos-Morales, F., Prieto, A. I., Beuzon, C. R., Holden, D. W. & Casadesus, J. Role for Salmonella enterica enterobacterial common antigen in bile resistance and virulence. J Bacteriol 185, 5328–5332, doi:10.1128/JB.185.17.5328-5332.2003 (2003).

27 Chng, S. S., Ruiz, N., Chimalakonda, G., Silhavy, T. J. & Kahne, D. Characterization of the two-protein complex in Escherichia coli responsible for lipopolysaccharide assembly at the outer membrane. Proceedings of the National Academy of Sciences of the United States of America 107, 5363–5368, doi:10.1073/pnas.0912872107 (2010).

28 Liu, D., Cole, R. A. & Reeves, P. R. An O-antigen processing function for Wzx (RfbX): a promising candidate for O-unit flippase. J Bacteriol 178, 2102–2107, doi:10.1128/jb.178.7.2102-2107.1996 (1996).

29 Sheng, H., Lim, J. Y., Watkins, M. K., Minnich, S. A. & Hovde, C. J. Characterization of an Escherichia coli O157:H7 O-antigen deletion mutant and effect of the deletion on bacterial persistence in the mouse intestine and colonization at the bovine terminal rectal mucosa. Appl Environ Microbiol 74, 5015–5022, doi:10.1128/AEM.00743-08 (2008).

30 Camprubi, S., Merino, S., Guillot, J. F. & Tomas, J. M. The role of the O-antigen lipopolysaccharide on the colonization in vivo of the germfree chicken gut by Klebsiella pneumoniae. Microb Pathog 14, 433–440, doi:10.1006/mpat.1993.1042 (1993).

31 Chaput, C., Spindler, E., Gill, R. T. & Zychlinsky, A. O-antigen protects gram-negative bacteria from histone killing. PloS one 8, e71097, doi:10.1371/journal.pone.0071097 (2013).

32 Miller, E. S. et al. Bacteriophage T4 genome. Microbiology and molecular biology reviews: MMBR 67, 86–156, table of contents, doi:10.1128/MMBR.67.1.86-156.2003 (2003).

33 Kutter, E. et al. Evolution of T4-related phages. Virus Genes 11, 285–297, doi:10.1007/BF01728666 (1995).

34 Thanassi, D. G., Cheng, L. W. & Nikaido, H. Active efflux of bile salts by Escherichia coli. J Bacteriol 179, 2512–2518, doi:10.1128/jb.179.8.2512-2518.1997 (1997).

35 Pazzi, P. et al. Bile salt-induced cytotoxicity and ursodeoxycholate cytoprotection: in-vitro study in perifused rat hepatocytes. Eur J Gastroenterol Hepatol 9, 703–709, doi:10.1097/00042737-199707000-00011 (1997).

36 Merritt, M. E. & Donaldson, J. R. Effect of bile salts on the DNA and membrane integrity of enteric bacteria. Journal of medical microbiology 58, 1533–1541, doi:10.1099/jmm.0.014092-0 (2009).

37 Cremers, C. M., Knoefler, D., Vitvitsky, V., Banerjee, R. & Jakob, U. Bile salts act as effective protein-unfolding agents and instigators of disulfide stress in vivo. Proceedings of the National Academy of Sciences of the United States of America 111, E1610–1619, doi:10.1073/pnas.1401941111 (2014).

38 Le, V. V. H., Biggs, P. J., Wheeler, D., Davies, I. G. & Rakonjac, J. Novel mechanisms of TolC-independent decreased bile-salt susceptibility in Escherichia coli. FEMS Microbiol Lett 367, doi:10.1093/femsle/fnaa083 (2020).

39 Perkins, H. R. Specificity of combination between mucopeptide precursors and vancomycin or ristocetin. Biochem J 111, 195–205, doi:10.1042/bj1110195 (1969).

40 Hernandez-Rocamora, V. M. et al. Coupling of polymerase and carrier lipid phosphatase prevents product inhibition in peptidoglycan synthesis. Cell Surf 2, 1–13, doi:10.1016/j.tcsw.2018.04.002 (2018).

41 Kanegasaki, S. & Wright, A. Mechanism of polymerization of the Salmonella O-antigen: utilization of lipid-linked intermediates. Proceedings of the National Academy of Sciences of the United States of America 67, 951–958, doi:10.1073/pnas.67.2.951 (1970).

42 Olivera, C., Le, V. V. H., Davenport, C. & Rakonjac, J. In vitro synergy of 5-nitrofurans, vancomycin and sodium deoxycholate against Gram-negative pathogens. Journal of medical microbiology 70, doi:10.1099/jmm.0.001304 (2021).

43 Ratib, N. R., Seidl, F., Ehrenreich, I. M. & Finkel, S. E. Evolution in Long-Term Stationary-Phase Batch Culture: Emergence of Divergent Escherichia coli Lineages over 1,200 Days. Mbio 12, doi:10.1128/mBio.03337-20 (2021).

44 Conter, A., Gangneux, C., Suzanne, M. & Gutierrez, C. Survival of Escherichia coli during long-term starvation: effects of aeration, NaCl, and the rpoS and osmC gene products. Res Microbiol 152, 17–26, doi:10.1016/s0923-2508(00)01164-5 (2001).

45 Lennox, E. S. Transduction of linked genetic characters of the host by bacteriophage P1. Virology 1, 190–206, doi:10.1016/0042-6822(55)90016-7 (1955).

46 Datsenko, K. A. & Wanner, B. L. One-step inactivation of chromosomal genes in Escherichia coli K-12 using PCR products. Proceedings of the National Academy of Sciences of the United States of America 97, 6640–6645, doi:10.1073/pnas.120163297 (2000).

47 Hong, Y. & Reeves, P. R. Diversity of o-antigen repeat unit structures can account for the substantial sequence variation of wzx translocases. J Bacteriol 196, 1713–1722, doi:10.1128/JB.01323-13 (2014).

48 Tran, E. N., Papadopoulos, M. & Morona, R. Relationship between O-antigen chain length and resistance to colicin E2 in *Shigella flexneri*. Microbiology 160, 589–601, doi:10.1099/mic.0.074955-0 (2014).

49 Qin, J. et al. A method for increasing electroporation competence of Gram-negative clinical isolates by polymyxin B nonapeptide. Scientific reports 12, 11629, doi:10.1038/s41598-022-15997-8 (2022).

50 CLSI. Methods for dilution antimicrobial susceptibility tests for bacteria that grow aerobically; approved standard. 11 edn, (Clinical and Laboratory Standards Institute, 2018).

