## Supplementary material for "O antigen biogenesis sensitises *Escherichia coli* K-12 to bile salts, a likely cause for how it lost its O antigen": SI

### Additional materials and methods for supplementary results

#### *RNase I leakage assay*

RNase I leakage assay was done as described previously[1].

#### *LPS silver staining*

LPS silver staining was done as described previously[2].

**Table S1. Strains, plasmids, and oligonucleotides**

**Bacterial strains**

| **Strains** | **Description** | **Source** |
|---|---|---|
| MG1655 | Wild-type *E. coli* K-12 MG1655 | Lab stock |
| MG1655*ΔwaaL* | MG1655*ΔwaaL::kan* | This work |
| MG1655*ΔtolC* | MG1655*ΔtolC::chl* | This work |
| MG1655-S | MG1655 with IS5I removed in *wbbL* | This work |
| MG1655-S*ΔrmlC* | MG1655-S*ΔrmlC::kan* | This work |
| MG1655-S*ΔwecA* | MG1655-S*ΔwecA::kan* | This work |
| MG1655-S*Δwzy* | MG1655-S*Δwzy::chl* | This work |
| MG1655-S*Δwzz* | MG1655-S*Δwzz::chl* | This work |
| MG1655-S*ΔwaaL* | MG1655-S*ΔwaaL::kan* | This work |
| MG1655-S*ΔwaaU* | MG1655-S*ΔwaaU::kan* | This work |
| BP1-BP51 | Library of 51 independent suppressor mutants of MG1655-S | This work |
| UT5600 | *E. coli* K-12, *ompT* | [3] |
| UT5600-S | UT5600 with IS5I removed in *wbbL* | This work |
| 2457T | *Shigella flexneri* 2a, 2457T | [4] |
| 2457T*ΔrmlD* | 2457T *ΔrmlD::kan* | [5] |
| EC958 | Uropathogenic *E. coli* cystitis isolate | [6] |
| EC958*ΔwaaL* | EC958*ΔwaaL::chl* | This work |
| EC958*ΔwecA* | EC958*ΔwecA::chl* | This work |
| CFT073 | Uropathogenic *E. coli* pyelonephritis isolate | [7] |
| CFT073*ΔwaaL* | CFT073*ΔwaaL::chl* | This work |
| TOP10 | F–mcrA Δ(mrr-hsdRMS-mcrBC) φ80lacZΔM15 ΔlacX74 recA1 araD139 Δ(ara-leu)7697 galU galK λ– rpsL(StrR) endA1 nupG | Invitrogen |

**Plasmids**

| **Plasmids** | **Description** | **Source** |
|---|---|---|
| pSU2718 | Cloning plasmid, lac promoter, $Chl^R$ | [8] |
| pBAD18cm | Cloning plasmid, arabinose promoter, $Chl^R$ | [9] |
| pKD46 | Temperature sensitive plasmid expressing Red proteins, $Amp^R$ | [10] |
| pKD4 | Plasmid carrying FRT flanked kanamycin resistant cassette, $Amp^R$, $Kan^R$ | [10] |
| pKD3 | Plasmid carrying FRT flanked kanamycin resistant cassette, $Amp^R$, $Chl^R$ | [10] |
| pWQ572 | Tetracycline inducible promoter, $Chl^R$ | [11] |
| pWbbL | *wbbL* CDS cloned from WG1 into pWQ572 | [12] |
| $pRmlC^{WT}$ | *rmlC* cloned from MG1655 into pBAD18cm | This work |
| $pRmlC^{L122W}$ | $rmlC^{L122W}$ cloned from BP27 into pBAD18cm | This work |
| pWaaL | *waaL* cloned from MG1655 into pSU2718 | This work |
| pJRD215 | Cosmid, $Kan^R$ | [13] |

| | |
|---|---|
| p*rfb*$^{sf}$ | pJRD215 carrying *rfb* region from *S. flexneri*, pRMA154 [14] |

**Oligos**

| **Description** | **Sequence** |
|---|---|
| *wbbL* CDS F/P1 | ATGGTATATATAATAATC |
| *wbbL* CDS R/P2 | TTATATTACGGGTGAAAA |
| *wbbL* screening F/P3 | ATTGACTTCAAAAAAGGTAACCTC |
| *wbbL* screening R/P4 | GAATGTTCGCAAATAAGTATACAAAG |
| *tolC* KO F | CCTTTTGCGGTAGCGGCTTCTGCTAGAATCCGCAATAATTTTACAGTGTAGGCTGGAGCTGCTTC |
| *tolC* KO R | CTCGCTGGCACCAACAAAGTTGTACTGGGCCTGTTTCACCATGGGAATTAGCCATGGTCC |
| *wzy* KO F | CGCTCTTTATCAAGTGAAAAATATAATGAGTACGGATTAAGTGTAGGCTGGAGCTGCTTC |
| *wzy* KO R | CGCGTCTAGAGAAATTTAAATCATTCAAAAAATACATTTTATGGGAATTAGCCATGGTCC |
| *wzz* KO F | AACTATGCGGACTTGGAAATTTCCGTCAGTTAGGGTAATGGTGTAGGCTGGAGCTGCTTC |
| *wzz* KO R | GGTGTCACCACCCTGCCCTTTTTCTTTAAAACCGAAAAGAATGGGAATTAGCCATGGTCC |
| K-12 *rmlC* KO F | ATGAATGTGATTAGAACTGAAATTGAAGATGTGCTAATTCTGGAGCCAAGGTGTAGGCTGGAGCTGCTTC |
| K-12 *rmlC* KO R | TCATGCAATTAATTTTAATCTGATAAGCTCATCTAACGTAAAGAGCCTTTATGGGAATTAGCCATGGTCC |
| *wecA* KO F | TCGGTTTACGCAGGGATTTGCTTCACGTTCGGAATTGTCGGTGTAGGCTGGAGCTGCTTC |
| *wecA* KO R | CTGCGTTTTACGCGCTTAATAAAGCGAGCAACTTTCCAGGATGGGAATTAGCCATGGTCC |
| *waaU* KO F | TTAATTAAGAATAAACAAGTTTAAGAAGTGAGTTAAAACATGGGAATTAGCCATGGTCC |
| *waaU* KO R | TAATTAATCATCCTGAAACTAAAATAATATGGTATAAAAGTGTAGGCTGGAGCTGCTTC |
| K-12 *waaL* KO F | TCAACAGTCAAGCAGTTTTGGAAAAGTTATCATCATTATAAAGGTAAAACATGGGAATTAGCCATGGTCC |
| K-12 *waaL* KO R | TTGTATAGATAAGAAGTGAGTTTTAACTCACTTCTTAAACTTGTTTATTCGTGTAGGCTGGAGCTGCTTC |
| EC958 *waaL* KO F | TCAACAGTCAAGCAGTTTTGGAAAAGTTATCATCATTATAAAGGTAAAACGTGTAGGCTGGAGCTGCTTC |

| | |
|---|---|
| EC958 *waaL* KO R | ATTAAGTTGTATAGATAAGAAGTGAGTTTTAACTCACTTCTTAAACTTGTATGGGAATTAGCCATGGTCC |
| CFT073 *waaL* KO F | GTTAGGTCTTGCATTAAATAACCCATCCTCGTAGCATAGGTTGAAATTATGTGTAGGCTGGAGCTGCTTC |
| CFT073 *waaL* KO R | TAATTTTGAAATAAAATCAGCTTCCTTGCTGATTTTATTTCACATATTCAATGGGAATTAGCCATGGTCC |
| K-12 *rmlC* cloning F | AATTTCTAGATCATGCAATTAATTTTAATCTGATAAGCTC |
| K-12 *rmlC* cloning R | AATTGGTACCATGAATGTGATTAGAACTGAAATTGAAG |
| K-12 *waaL* cloning F | AATTGGTACCATGCTAACATCCTTTAAACTTCATTC |
| K-12 *waaL* cloning R | AATTGGATCCTTAATTAATTGTATTGTTACGATTATTAATGACG |

**Table S2. Differences of MG1655 used in this study and reference MG1655 (U00096)**

| **Parent MG1655** | | |
|---|---|---|
| **Mutations** | **locus in U00096** | **Note** |
| Deletion of Insertion element between *frsA* and *crl* | 257927-258623 | |
| Deletion of mobile element between *ychE* and *oppA* | 1299498-1300694 | |
| Deletion between *ynaJ* and *dbpA* | 1397459-1411026 | |
| Deletion of mobile element between *flhD* and *uspC* | 1978516-1979229 | |
| Deletion of 2 bases (CC) in *gatC* | 2173363-2173364 | Frame shift |
| Single base (G to T) substitution in *ypjB* | 2784096 | |
| Single base (G) insertion in *glpG* | 3560455-3560456 | Frame shift |
| Double base insertion (GC) in REP321j region | 4296381-4296382 | |
| **Heterogeneous mutations in MG1655-S and Suppressor Mutants** | | |
| **Mutations** | **locus in U00096** | **Note** |
| Single base (A to T) substitution in *wbbL* (MG1655-S, BP2, BP5, BP11, BP27, BP28, BP30) | 2101747 | synonymous |
| Inversion between *ycfK* and *stfE* (BP16, BP27, BP28, BP30, | 1207790-1207805 | e14 prophage phase variation[15] |

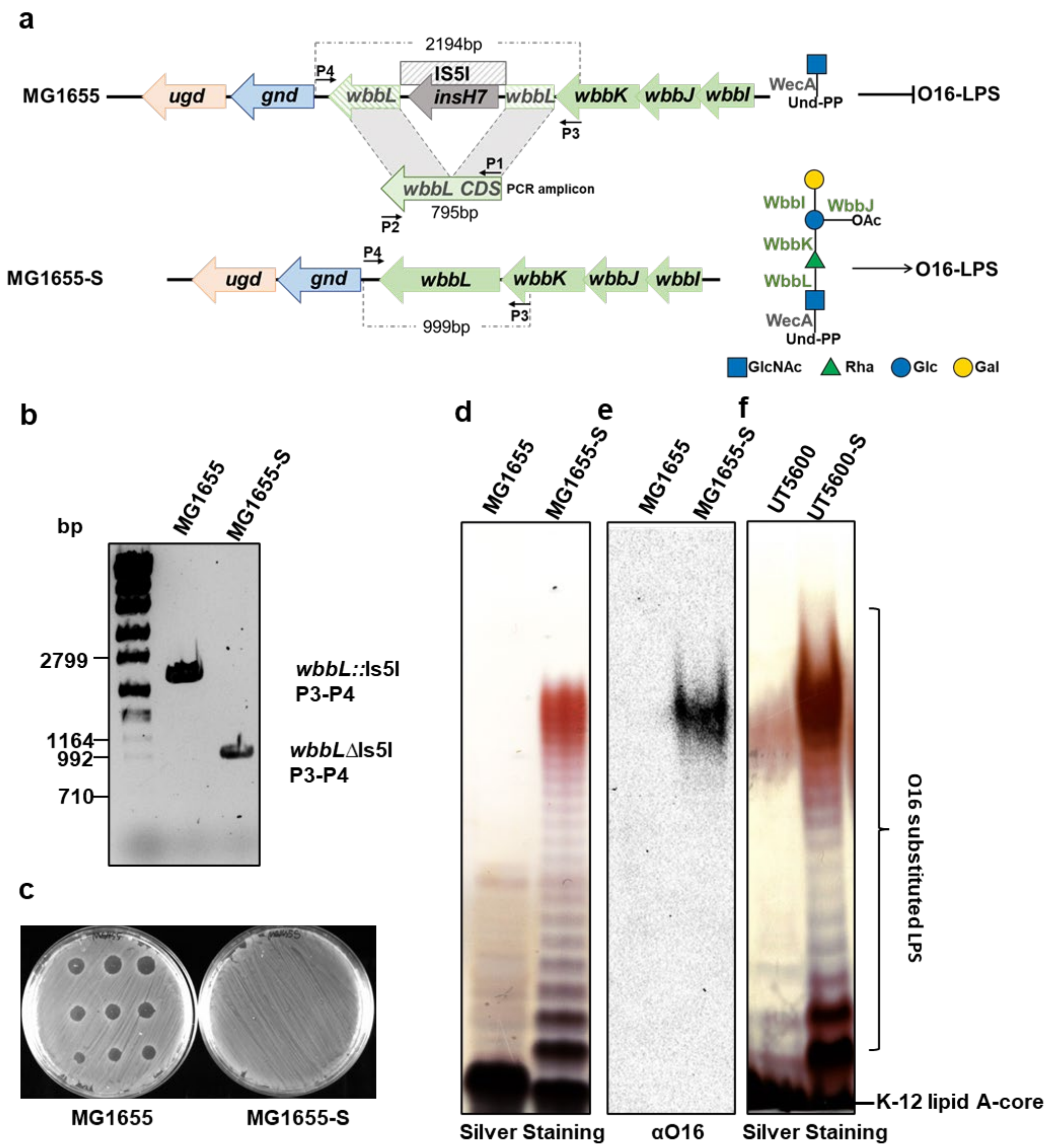


**Fig S1. Restoration of O16 production in *E. coli* K-12 strains. (a)** Schematic representation of the strategy employed for *wbbL* allelic replacement in the *rfb* gene cluster in MG1655 to construct strain MG1655-S. The respective O16 repeating units that would be produced in the cytosol denote the expected LPS products by each strain. Primers P1 and P2 used to generate MG1655-S are mapped onto *rfb* regions by black arrows. **(b)** PCR amplicons of *wbbL* region from MG1655 and MG1655-S (using primers P3 and P4) confirming the replacement of an intact *wbbL* CDS in MG1655-S. **(c)** Colicin sensitivity assay confirming increased resistance of MG1655-S due to regained O antigen substituted LPS production. Colicin E2 was used at 1 mg/ml and in subsequent 2-fold dilutions (5 µl spots). (**d&f**) Silver stained SDS-PAGE of LPS samples from two sets of *E. coli* K-12 strains (wild-type MG1655 and UT5600 carry the IS51 element in *wbbL* and MG1655-S/UT5600-S are isogenic *wbbL* intact strains, respectively). LPS patterns confirm restored production of O antigen substituted LPS in the engineered MG1655-S and UT5600-S strains. (**e**) Western immunoblotting of samples as in (**d**) with anti-O16 antibodies showing the restoration of O16 O antigen production in MG1655-S.

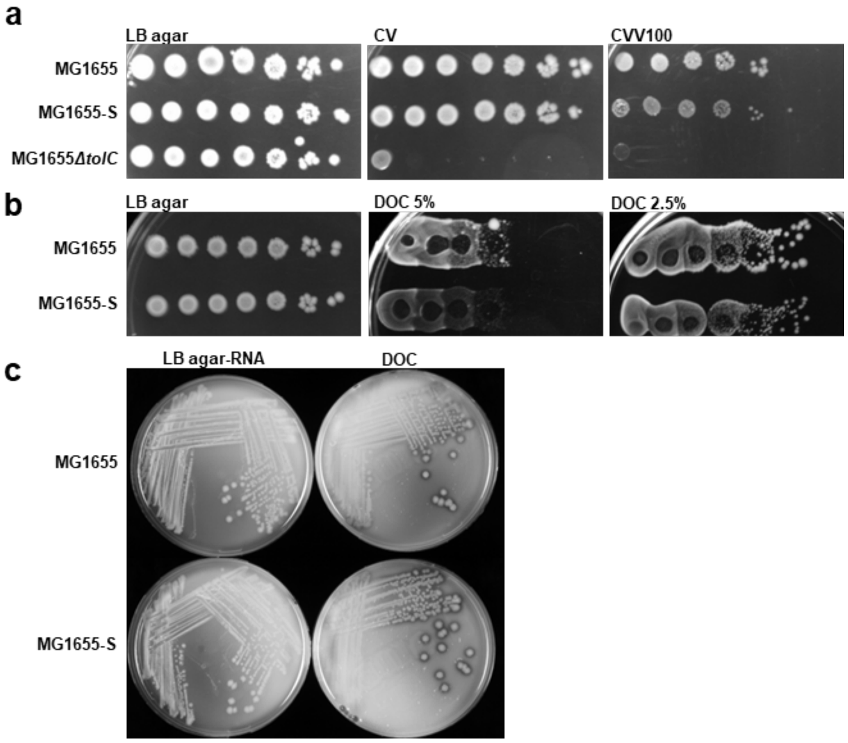


**Fig S2. The effect of crystal violet and DOC on *E. coli* K-12 MG1655 and MG1655-S. (a&b)** Bacterial cultures of indicated strains grown in LB media were adjusted to OD600 of 1 and spotted (4 µl) in 10-fold serial dilutions ($10^0$ to $10^{-6}$) onto LB agar supplemented with 0.0001% (w/v) crystal violet (CV) or 100 µg/ml vancomycin and CV (CVV100), or 5% and 2.5% (w/v) sodium deoxycholate (DOC). (**c**) RNase I leakage assay of MG1655 and MG1655-S grown on LB agar supplemented without or with 0.1% (w/v) DOC.

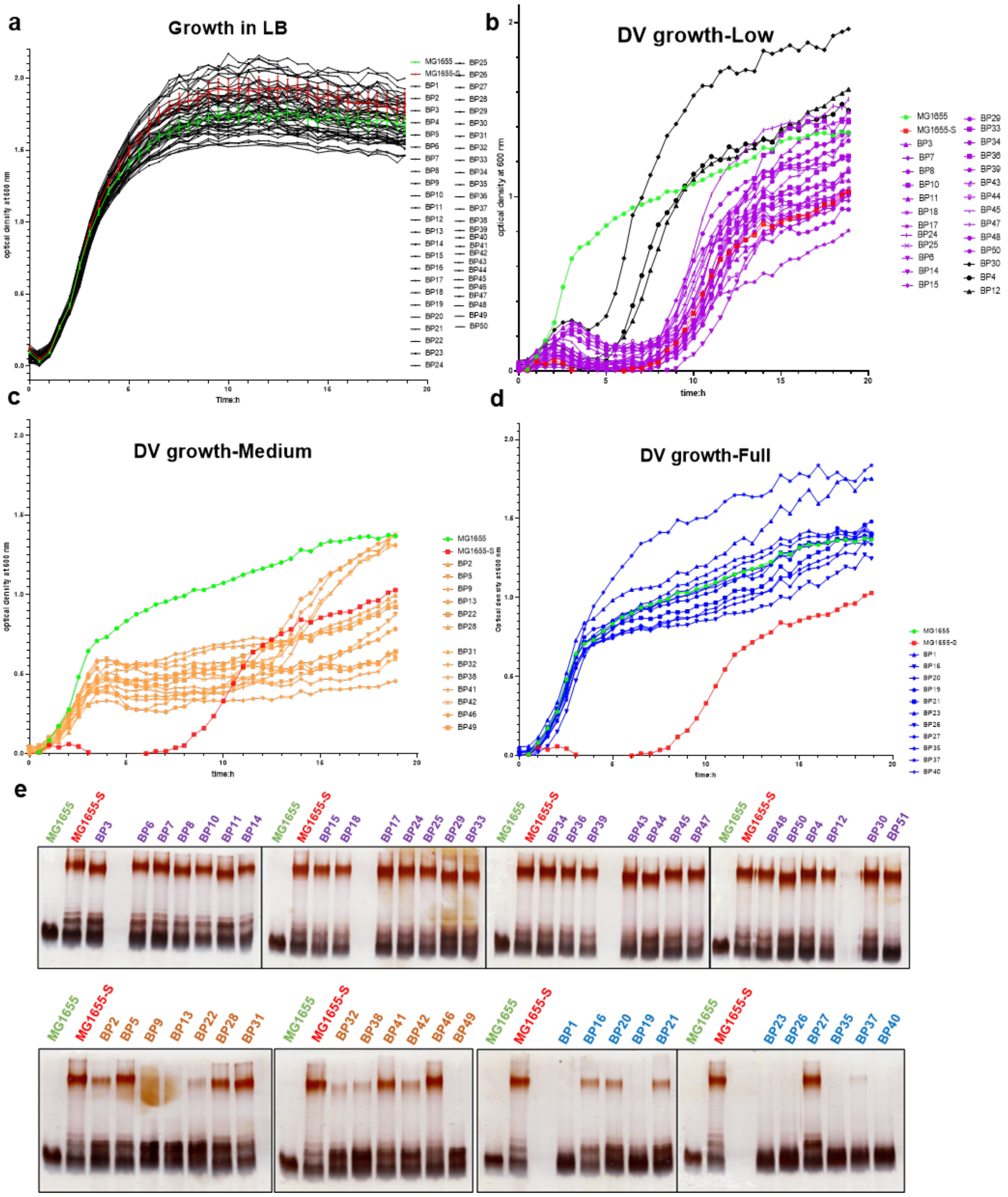


**Fig S3. Phenotypic analyses of 51 MG1655-S suppressor mutants.** Growth curve of MG1655, MG1655-S and BP1-BP51 suppressor mutants in LB media (**a**) or in LB supplemented with (**b-d**) 100 µg/ml vancomycin and 0.1% (w/v) DOC (DV100). **e**) Silver staining of SDS-PAGE of LPS samples prepared from MG1655, MG1655-S and BP1-BP51 suppressor mutants.

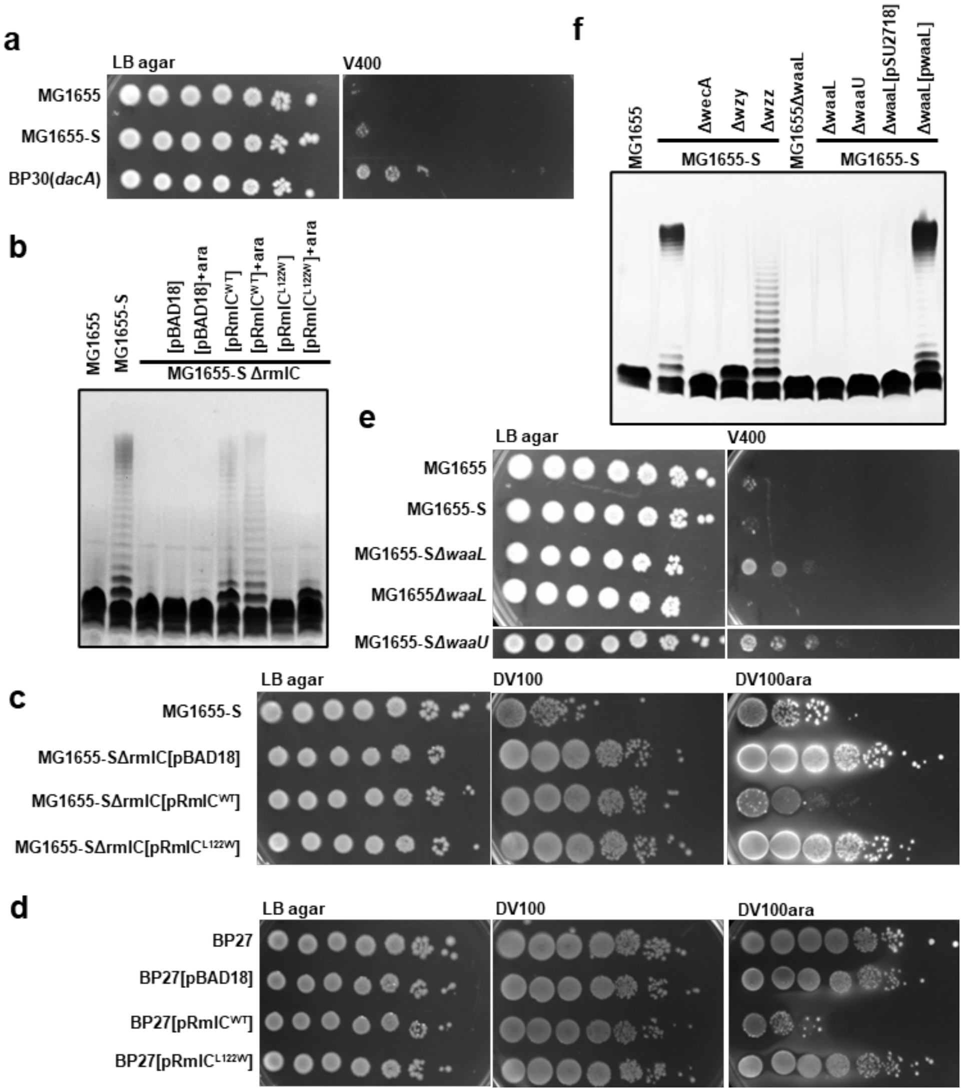


**Fig S4. Effect of DOC and vancomycin on MG1655-S mutants.** Bacterial cultures of indicated strains (**a, c-e**) grown in LB media were adjusted to OD600 of 1 and spotted (4 µl) in 10-fold serial dilutions ($10^0$ to $10^{-6}$) onto LB agar supplemented without or with 400 µg/ml vancomycin (V400), or 0.1% (w/v) DOC and 100 µg/ml vancomycin (DV100) in the absence or presence of 0.2% (w/v) L-arabinose (DV100ara). (**b & f**) Silver staining of SDS-PAGE of LPS samples of MG1655, MG1655-S and its suppressor mutants. Supplementation of media with 0.2% (w/v) arabinose was used in samples as indicated (+ara).

### References to supplementary information


1 Lopes, J., Gottfried, S. & Rothfield, L. Leakage of periplasmic enzymes by mutants of Escherichia coli and Salmonella typhimurium: isolation of "periplasmic leaky" mutants. *J Bacteriol* **109**, 520-525, doi:10.1128/jb.109.2.520-525.1972 (1972).

2 Qin, J. *Investigation of the IcsA-mediated Shigella flexneri hyper-adherence* PhD in Biological Sciences thesis, The University of Adelaide, (2022).

3 McIntosh, M. A., Chenault, S. S. & Earhart, C. F. Genetic and physiological studies on the relationship between colicin B resistance and ferrienterochelin uptake in Escherichia coli K-12. *J Bacteriol* **137**, 653-657, doi:10.1128/jb.137.1.653-657.1979 (1979).

4 Qin, J., Doyle, M. T., Tran, E. N. H. & Morona, R. The virulence domain of Shigella IcsA contains a subregion with specific host cell adhesion function. *PloS one* **15**, e0227425, doi:10.1371/journal.pone.0227425 (2020).

5 Tran, E. N., Doyle, M. T. & Morona, R. LPS unmasking of *Shigella flexneri* reveals preferential localisation of tagged outer membrane protease IcsP to septa and new poles. *PloS one* **8**, e70508, doi:10.1371/journal.pone.0070508 (2013).

6 Totsika, M. *et al.* Insights into a multidrug resistant Escherichia coli pathogen of the globally disseminated ST131 lineage: genome analysis and virulence mechanisms. *PloS one* **6**, e26578, doi:10.1371/journal.pone.0026578 (2011).

7 Welch, R. A. *et al.* Extensive mosaic structure revealed by the complete genome sequence of uropathogenic Escherichia coli. *Proceedings of the National Academy of Sciences of the United States of America* **99**, 17020-17024, doi:10.1073/pnas.252529799 (2002).

8 Martinez, E., Bartolome, B. & de la Cruz, F. pACYC184-derived cloning vectors containing the multiple cloning site and lacZ alpha reporter gene of pUC8/9 and pUC18/19 plasmids. *Gene* **68**, 159-162, doi:10.1016/0378-1119(88)90608-7 (1988).

9 Guzman, L. M., Belin, D., Carson, M. J. & Beckwith, J. Tight regulation, modulation, and high-level expression by vectors containing the arabinose PBAD promoter. *J Bacteriol* **177**, 4121-4130, doi:10.1128/jb.177.14.4121-4130.1995 (1995).

10 Datsenko, K. A. & Wanner, B. L. One-step inactivation of chromosomal genes in Escherichia coli K-12 using PCR products. *Proceedings of the National Academy of Sciences of the United States of America* **97**, 6640-6645, doi:DOI 10.1073/pnas.120163297 (2000).

11 Larue, K., Ford, R. C., Willis, L. M. & Whitfield, C. Functional and structural characterization of polysaccharide co-polymerase proteins required for polymer export in ATP-binding cassette transporter-dependent capsule biosynthesis pathways. *The Journal of biological chemistry* **286**, 16658-16668, doi:10.1074/jbc.M111.228221 (2011).

12 Hong, Y. & Reeves, P. R. Diversity of o-antigen repeat unit structures can account for the substantial sequence variation of wzx translocases. *J Bacteriol* **196**, 1713-1722, doi:10.1128/JB.01323-13 (2014).

13 Davison, J., Heusterspreute, M., Chevalier, N., Ha-Thi, V. & Brunel, F. Vectors with restriction site banks. V. pJRD215, a wide-host-range cosmid vector with multiple cloning sites. *Gene* **51**, 275-280, doi:10.1016/0378-1119(87)90316-7 (1987).

14 Morona, R., Mavris, M., Fallarino, A. & Manning, P. A. Characterization of the rfc region of Shigella flexneri. *J Bacteriol* **176**, 733-747, doi:10.1128/jb.176.3.733-747.1994 (1994).

15 Goldberg, A., Fridman, O., Ronin, I. & Balaban, N. Q. Systematic identification and quantification of phase variation in commensal and pathogenic Escherichia coli. *Genome Med* **6**, 112, doi:10.1186/s13073-014-0112-4 (2014).
